# Multi-omics analysis of longitudinal patient samples reveals the molecular mechanism of AML progression

**DOI:** 10.1101/2024.05.28.593678

**Authors:** Nisar Ahmed, Irene Cavattoni, William Villiers, Chiara Cugno, Sara Deola, Borbala Mifsud

## Abstract

Relapse remains a determinant of treatment failure and contributes significantly to mortality in acute myeloid leukemia (AML) patients. Despite efforts to understand AML progression and relapse mechanisms, findings on acquired gene mutations in relapse vary, suggesting inherent genetic heterogeneity and emphasizing the role of epigenetic modifications. We conducted a multi-omics analysis using Omni-C, ATAC-seq, and RNA-seq on longitudinal samples from two adult AML patients at diagnosis and relapse. Herein, we characterized genetic and epigenetic changes in AML progression to elucidate the underlying mechanisms of relapse. Differential interaction analysis showed significant 3D chromatin landscape reorganization between relapse and diagnosis samples. Comparing global open chromatin profiles revealed that relapse samples had significantly fewer accessible chromatin regions than diagnosis samples. In addition, we discovered that relapse-related upregulation was achieved either by forming new active enhancer contacts or by losing interactions with poised enhancers/potential silencers. Altogether, our study highlights the impact of genetic and epigenetic changes on AML progression, underlining the importance of multi-omics approaches in understanding disease relapse mechanisms and guiding potential therapeutic interventions.

## Introduction

Acute myeloid leukemia (AML) manifests as a complex disease marked by a multitude of genetic mutations and dysregulated gene expression profiles stemming from genetic and epigenetic alterations. These factors shape the trajectory of AML progression and confer resistance to therapeutic modalities. In general, AML occurs at any age, but it is the most prevalent form of acute leukemia in adults with a median age at diagnosis of 68 years and an estimated 20,380 diagnoses and 11,310 related deaths projected for 2023 (Kishtagari & Levine, 2021; Siegel et al., 2023). In the past decade, extensive research has focused on AML heterogeneity at disease onset, leading to improved classification (Döhner et al., 2022) and novel treatment agents (DiNardo & Cortes, 2016; Stein et al., 2017; Stone et al., 2017) that considerably help to achieve complete remission in most patients, however, the 5-year overall survival (OS) rates are still only at around 28% (Howlader N, et al., 2021). It is mainly due to the high relapse rate, as 40% to 60% of patients relapse within 3 years and fail to respond to conventional chemotherapy regimen (Bejanyan et al., 2015; Karlsson et al., 2017; Verma et al., 2010). Unfortunately, most individuals who relapse ultimately die from the disease (Schlenk et al., 2018). Prognosis in case of relapse is typically more unfavorable, especially when the recurrence occurs within a year after the initial remission (Rasche et al., 2021). Furthermore, relapse is a primary factor contributing to treatment failure (Döhner et al., 2015).

Recently, several investigations have leveraged next-generation sequencing (NGS) to identify gene mutations specific to relapse in certain AML subgroups, aiming to elucidate the disease’s course. However, there is considerable genetic heterogeneity among different relapsed patients (Jan et al., 2012; Metzeler et al., 2016; Papaemmanuil et al., 2016), even when focusing on specific AML subgroups (Ahn et al., 2018; Gröschel et al., 2015; Wang et al., 2016). This diversity hinders the discovery of consistent relapse-specific signatures. Additionally, there are only a few studies that compare the mutational profiles of diagnosis and relapse samples in AML patients, and they focus on coding mutations (Farrar et al., 2016; Masetti et al., 2016), despite growing evidence that non-coding mutations in regulatory elements or structural variants altering enhancer usage can also drive oncogenesis (Northcott et al., 2014; Zhu et al., 2020). This underscores the significance of epigenetic changes in the course of the disease and emphasizes the necessity of considering the intrinsic and extrinsic disease heterogeneity (Levin et al., 2021; Schwenger & Steidl, 2021; Vicente-Dueñas et al., 2018). Furthermore, there is a paucity of comprehensive molecular characterization of longitudinal AML samples, including diagnosis and relapse pairs. Therefore, we aim to assess the contribution of epigenetic factors, such as active regulatory elements and their long-range chromatin interactions, in conjunction with gene expression profiles across longitudinal samples, elucidating their role in the progression of AML.

To achieve this, we have profiled the long-range chromatin interactions (using Omni-C), open chromatin regions (using ATAC-seq), and gene expression (using RNA-seq) of matched adult AML patients at diagnosis and relapse. We found significant alterations in the 3D chromatin landscape and accessible chromatin regions between relapse and diagnosis samples.

Additionally, upregulated genes in relapse showed enrichment for H3K27me3 in distal regions of diagnosis-specific interactions, indicating loss of potential silencer connections during relapse.

## Results

### Overview of multi-omics assays in diagnosis and relapse AML

We integrated changes of the 3D genome structure, chromatin accessibility, and gene expression to decipher the molecular changes that occur in AML at relapse compared to at the time of diagnosis in two initial diagnoses and relapse AML sample pairs (Table 1). We explored long-range regulatory interaction patterns using Omni-C, active regulatory elements using ATAC-seq, and transcriptional patterns through RNA-seq (Figure 1A). In addition, we used Omni-C data to create genome-wide contact maps and assess the spectrum of large chromosomal changes in these AML samples. The contact maps identified that all of the samples had a t(9:11) or *KMT2A::MLLT3* translocation. This confirmed the initial diagnosis of one patient; however, this translocation was undetected by cytogenetics in the other patient. The KMT2A::MLLT3 subtype of AML is shown to have a poor/intermediate prognosis, and the current mechanistic understanding of KMT2A-rearrangement (KMT2Ar) prognosis has not fully translated into therapeutic success due to the complexity of genomic events contributing to the disease (Krivtsov & Armstrong, 2007; Liedtke & Cleary, 2009; Meyer et al., 2018). We additionally observed within our Omni-C data that relapse samples had gained extra chromosomal abnormalities, for example, chromosome 8 duplication, t(3:5), and t(1:X) (Figure 1B). Such abnormalities are common occurrences at relapse in AML (Kern et al., 2002).

**Figure 1.**
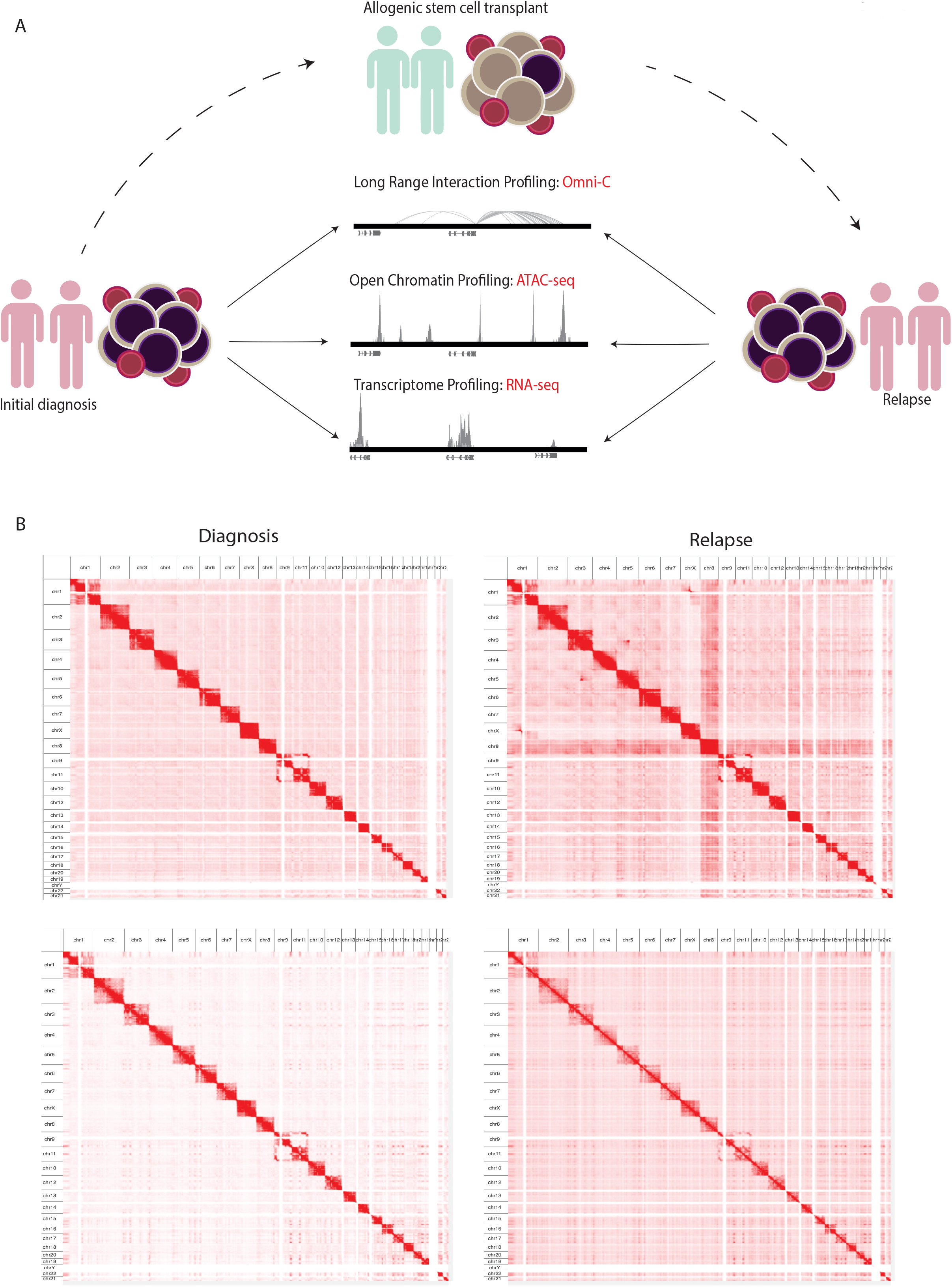
Study schematic showing multi-omics approaches in analyzing adult AML sample pairs. **A)** Schematic shows the experimental workflow where PBMCs obtained from adult AML samples at initial diagnosis undergo NGS library preparation for Omni-C, ATAC-seq, and RNA- seq analyses. Subsequently, the same NGS libraries were prepared from samples upon relapse following allotx. **B)** Contact maps derived from the Omni-C dataset, depicting the chromosomal architecture of diagnosis samples (*left*) compared with relapse samples (*right*).

**Table 1.** Clinical characteristics of AML patients at diagnosis and relapse.

| Serial # | Sample ID | Sample Type | Blasts (%) | Karyotype |
| --- | --- | --- | --- | --- |
| 1 | TMD | Diagnosis | 61% | 46, XX |
| 2 | TMR | Relapse | 88% | Complex |
| 3 | MBD | Diagnosis | >50% | NA |
| 4 | MBR | Relapse | >50% | NA |

### Differential interactions and differentially accessible regions in relapse versus diagnosis samples

Next, we asked whether there are any regions that are differentially interacting in the genome distinguishing relapse from diagnosis samples. While we found higher intra-patient similarity than intra-status (Figure 2A), we could detect common diagnosis and relapse-specific interactions. By comparing relapse versus diagnosis samples, we found 8,202 significant differential interactions (p-adj < 0.1), where 5,262 interactions were defined as diagnosis-specific, and 2,940 interactions were found to be relapse-specific (Figure 2B). This highlights there are distinctive interaction patterns between diagnosis and relapse timepoints that are consistent across patients. Next, we explored the chromatin accessibility profile of the diagnosis and relapse pairs using ATAC-seq. We estimated the similarity between relapse and diagnosis samples using PCA based on the global open chromatin profile, which again shows higher inter-patient variability than inter-state variability (Figure 2C). Subsequently, we identified 16,363 consistent diagnosis-specific open chromatin regions and 6,966 relapse-specific regions in addition to 4,099 regions that were active in all samples (Figure 2D).

**Figure 2.**
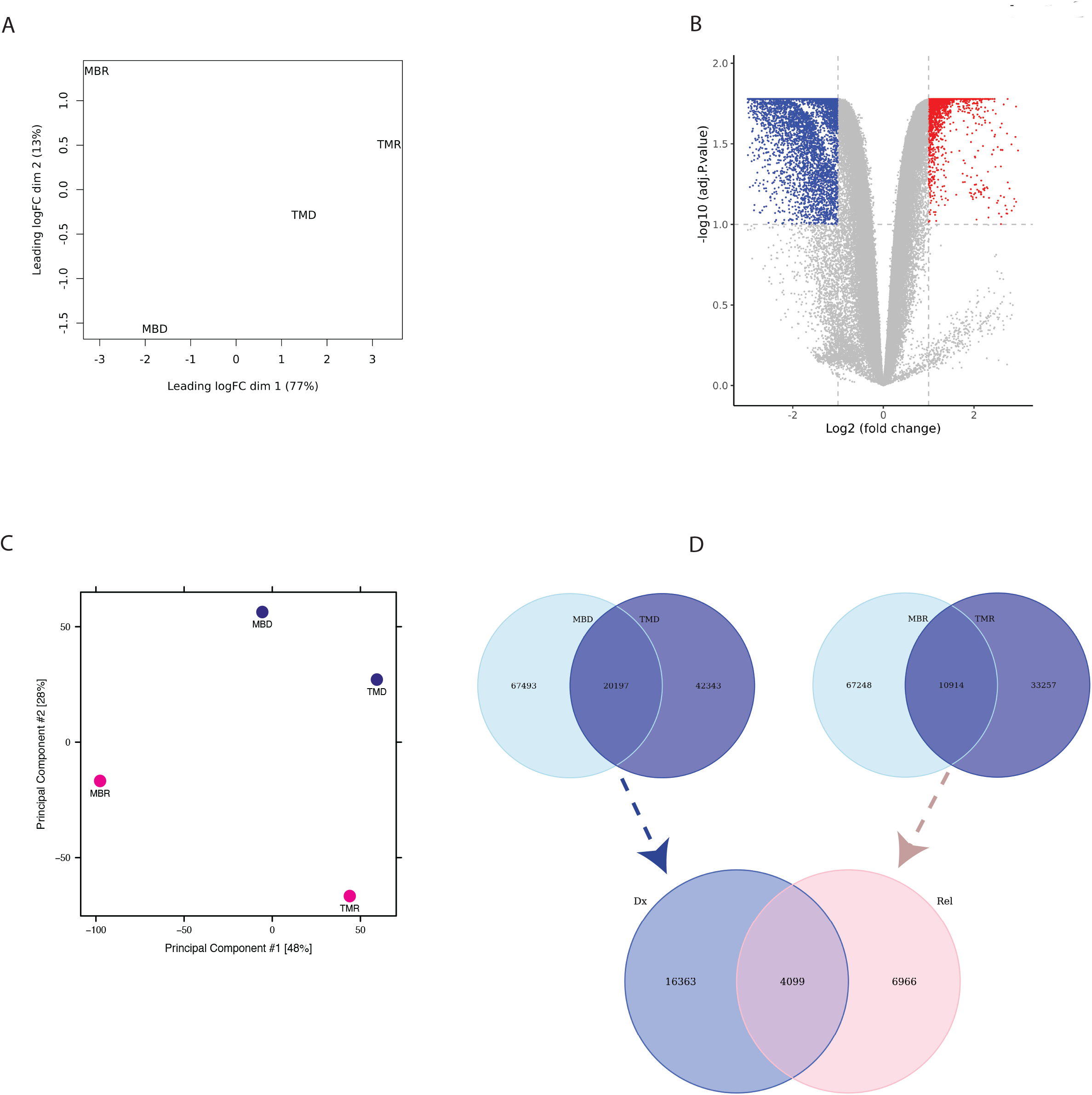
Illustrating the differential interactions analysis and overlap of ATAC-seq peaks. **A)** PCA plot based on limma logFC of Omni-C data showing higher inter-patient similarity **B)** Volcano-plot showing differential interaction of relapse versus diagnosis based on Omni-C datasets where each dot represents an interaction. The blue represents significant diagnosis-specific interactions (5,262), and red represents significant relapse-specific interactions (2,940), whereas grey dots represent none significant interactions. **C)** Similar PCA plot, but here it is based on ATAC-seq normalized peaks showing inter-status similarity where diagnosis samples have similar chromatin accessibility profiles. **D)** Venn diagram representing the overlap of diagnosis-specific ATAC-seq peaks (*upper left*), relapse-specific ATAC-seq peaks (*upper right*), and diagnosis-specific and relapse-specific ATAC-seq peaks (*bottom*).

### Integration of differential chromatin interaction and accessibility signatures

We integrated the Omni-C anchors with ATAC-seq peaks to investigate which differential interacting regions are also differentially accessible (Supplementary Figure 1A). We found 150 unique relapse-specific anchors that were differentially accessible in relapse samples, which were annotated with 107 genes. GO-term analysis of these genes reveals the biological process and pathways in which these genes were enriched, such as regulation of canonical Wnt signaling pathway, negative regulation of cell differentiation, Notch signaling pathway, and AML (Figure 3A). Next, we investigated if relapse-specific peaks in relapse-specific anchors are associated with different TFs compared to diagnosis-specific ones by performing differential motif discovery on these two sets. We noted that a liver X receptor beta (LXRb) was enriched in relapse-specific regions (Figure 3B). Finally, we explored whether these differential interactions are anchored at active or poised enhancers. For instance, *HOXA9*, which is involved in cell differentiation and upon dysregulation, contributes to leukemogenesis, was found to form a relapse-specific long-range interaction with an active enhancer at the 3’UTR of the *SNX10* gene, marked by H3K4me1/3 and H3K27ac (Figure 3C). In addition, we found KMT2Arr subtype-specific patterns at known KMT2A target genes, such as *UBE2J1* and *PARP8*, in the loop anchors (Supplementary Figure 1B,C). Overexpression of these genes contributes to the blockage of normal hematopoietic differentiation and promotes leukemogenesis.

**Figure 3.**
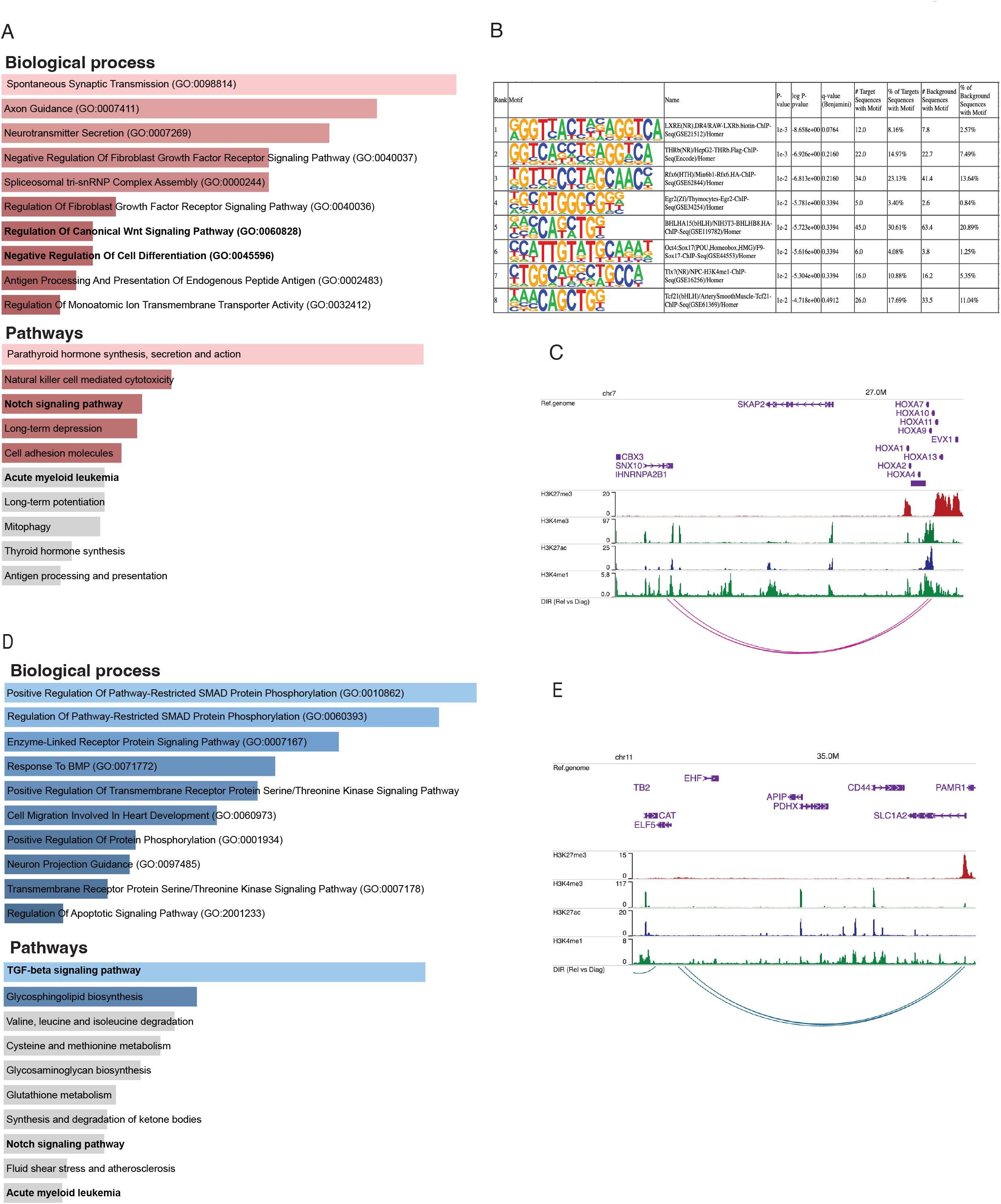
Integration analysis of Omni-C interaction and ATAC-seq peaks. **A)** Bar plots representing the enrichment of biological processes and pathways based on genes annotated to those regions where both interactions and peaks overlapped in relapse samples. **B)** The top 8 motifs enriched in relapse-specific Omni-C anchors and ATAC-seq peaks compared to remission anchors and peaks. **C)** Arc plot from WashU Epigenome browser representing relapse-specific Omni-C interaction and ATAC-seq peaks where *HOXA9* region interacts with distant enhancer at the downstream region of the *SNX10* gene. The enhancer was validated using publicly available annotation data in the browser, where it was enriched for H3K4me3/1 and H3K27ac but lacked enrichment for H3K27me3, suggesting an active enhancer. **D)** Similar bar plots like in **A**, but here the genes are annotated from diagnosis-specific interactions and peaks. **E)** Similar Arch plot like in **C**, but here showing diagnosis-specific interaction and peaks at the promoter region of *ELF5* interacting with poised enhancer, which is marked H3K27me3 and H3K4me1.

On the other hand, we found 312 unique diagnosis-specific anchors that were differentially accessible and in proximity of these peaks, 154 genes were present. Further, these genes were enriched in biological pathways such as TGF-beta and Notch signaling (Figure 3D). Examples of genes with diagnosis-specific interactions include *ELF5*, which is associated with epithelial cell function and has implications in cancer. Its promoter formed a diagnosis-specific contact with a poised enhancer marked by H3K27me3 and H3K4me1, contributing to its low expression in diagnosis samples (Figure 3E). Additionally, a TGF-beta pathway member, *BMP8A*, also interacted with a poised enhancer. This gene plays critical roles in various cellular processes, such as tissue differentiation, cell proliferation, and apoptosis. *ST8SIA1*, which is involved in cell signaling, recognition, and adhesion, was interacting with an active enhancer near the 3’UTR of *CMAS*. Another gene with a diagnosis-specific active enhancer contact is *HPSE2*, which has a in tumor microenvironment dynamics and cancer progression. Its promoter is interacting with an active enhancer present in the proximity of its own 3’UTR (Supplementary Figure 1D-F).

### Analysis of differential gene expression dynamics

Finally, we explored the gene expression profile of these longitudinal samples using RNA-seq. Similarly to what we found using the other omics methods, samples do not cluster together based on disease status i.e., relapse and diagnosis (Figure 4A). Nonetheless, we asked whether there are differentially expressed genes between relapse versus diagnosis samples. Using limma-voom, we found overall, 95 significantly dysregulated genes that were selected using p.adj < 0.1 and absolute log2FC > 1 (Figure 4B). These genes were enriched in pathways like B-cell receptor signaling pathway and immune and defense mechanism-related biological processes as well as AML (Supplementary Figure 3A). Further, out of 95 significant dysregulated genes, there were 55 upregulated genes (Supplementary Table 1) and 40 downregulated genes (Supplementary Table 5.2). These upregulated genes were enriched in defense response and interferon and cytokines signaling (Figure 4C), whereas downregulated genes were enriched in B-cell receptor, Jak-STAT, and interleukin signaling pathways (Figure 4D).

**Figure 4.**
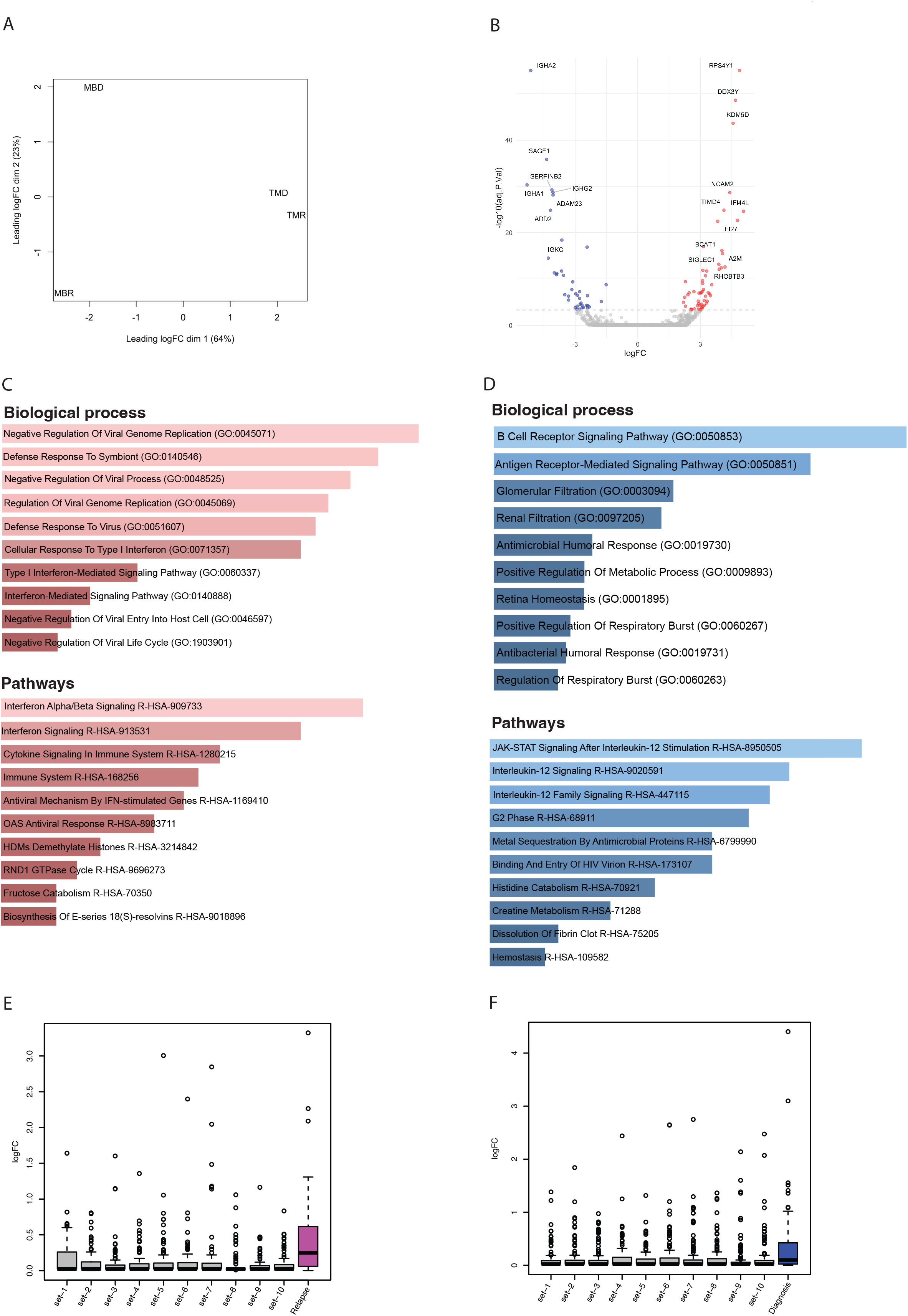
Differential gene expression analysis and integration with Omni-C and ATAC- seq. **A)** PCA plot showing higher inter-patient variability based logFC instead of inter-status variability. **B)** Volcano plot showing significant (p-adj < 0.1) differentially expressed genes from limma; here, blue dots are significantly down-regulated genes, red dots are significantly up- regulated, and grey are none significant genes. **C)** Bar plots showing the enrichment of biological processes and pathways based on up-regulated genes. **D)** Similar bar plots as **C,** but here, it is showing enrichment of biological processes and pathways based on down-regulated genes. **E)** Box plots representing up-regulated genes that overlapped with relapse-specific interactions and peaks and their comparison with the expression level of 10 different random sets of up-regulated genes. **F)** Similar to the Box plots in **E** but here the up-regulated genes overlapped with diagnosis-specific interactions and peaks and its comparison with expression level of 10 different random sets of up-regulated genes.

### Multi-omics data integration exploration

Finally, we integrated all three omic datasets together to identify consistently differentially regulated genes across relapse and diagnostic timepoints. While there were only a handful of genes that showed significant differences, we noted that in general, genes that were associated with relapse-specific interactions and open chromatin regions showed higher levels of upregulation than those genes associated with diagnosis-specific interactions and open chromatin regions, as well as random genes from the genome, indicating that relapse-specific interactions are often new active enhancer contacts (Figure 4E,F). Those genes that were associated with diagnosis-specific interactions and open chromatin regions showed slightly higher levels of upregulation than random genes from the genome (Figure 4F). We hypothesized that these up- regulated genes may have initially interacted with silencers/poised enhancers, but upon relapse, this interaction is lost (Supplementary Figure 2B). We found 13 genes (*ST8SIA1*, *IVD*, *ACAN*, *KIF5C*, *ADAMTS5*, *CFAP299*, *PITX2*, *KLHL31*, *RBPMS*, *SNAI2*, *SOX17*, *CSMD3*, and *DMRTA1*) where the diagnosis-specific interaction linked them to regions marked by H3K27me3, which is a significantly higher proportion than expected by chance (p-value=1.016x10^-05^). In contrast, genes associated with either relapse-specific or diagnosis-specific interactions and open chromatin regions showed a similar level of downregulation compared to random genes from the genome (Supplementary Figure 2C,D).

There were five genes that showed significant differences in all three analyses. Our findings revealed that among genes that were upregulated in relapse, *SDC2* and *CD70* were in consistent relapse-specific anchors and peaks, while *NCAM2* and *IFI44* were in consistent diagnosis-specific anchors and peaks. These genes have been shown to have some link with different types of cancers. For example, the SDC2 (syndecan-2) protein functions as an integral membrane protein and participates in cell proliferation, cell migration and cell-matrix interactions via its receptor for extracellular matrix proteins, and altered *SDC2* expression has been detected in several different tumor types (Akl et al., 2015; Canarte et al., 2023). Among the down-regulated genes in relapse, *TSPYL5*, which is a *TP53* suppressor via its interaction with *USP7* (Epping et al., 2011), was found in a consistent diagnosis-specific anchor and peak. We explored the relationship between the expression of these genes and clinical outcome, in terms of survival, using the Leucegene AML RNA-seq prognostic cohort (n= 373). We observed that the upregulation of these genes, including *TSPYL5*, negatively impacts the overall survival of AML patients, except for *CD70*, which did not show a significant effect (Supplementary Figure 3A-E).

## Discussion

AML aggressive subtypes are often linked to rearrangements in the mixed lineage leukemia (KMT2Ar) gene. 10% of adult acute leukemias with a very poor prognosis and chemoresistance are caused by clinically significant and genetically well-defined KMT2Ar leukemia (Krivtsov & Armstrong, 2007; Liedtke & Cleary, 2009; Meyer et al., 2018). Additionally, patients with t(9;11)(p22;q23), the most frequent translocation which leads to the *KMT2A::MLLT3* fusion gene, show relatively acceptable results with intensive chemotherapy (Grimwade et al. 2010; Mrózek et al. 1997; Stölzel et al. 2016; Chen et al. 2013; Pigneux et al. 2015), placing them in the intermediate risk group according to ELN 2017 and ELN 2022 classification (Döhner et al., 2017, 2022). Our chromatin conformation data revealed an undetected 9;11 translocation in one patient. This highlights the need for more in-depth karyotyping by techniques like those using next generation sequencing. Techniques such as targeted RNA-seq are starting to become routine in such diagnosis to detect low-level fusion genes (Kerbs et al., 2022).

Advances in pharmacological inhibitors and targeted immuno-therapies have considerably improved the treatment options for KMT2Ar leukemias (Issa et al., 2023; van der Sluis et al., 2023). However, KMT2Ar leukemias show highly heterogenous responses to therapeutic regimens despite their similar oncogenic lesions (Issa et al., 2023; Lambo et al., 2023). Common genetic mechanisms leading to therapeutic resistance include clonal selection and the acquisition of secondary mutations. Additionally, a subset of these leukemias may also evade targeted therapies through epigenetic mechanisms that remain poorly understood (Tirtakusuma et al., 2022).

In this study, we aimed to elucidate the role of epigenetic modifications and 3D genome organization in AML progression by integrating ATAC-seq and RNA-seq data with high-resolution chromosome conformation capture (Omni-C) in patient samples obtained at diagnosis and relapse stages. Our findings revealed differences in the 3D chromatin architecture between relapse and diagnosis samples, characterized by a general loss of significant chromatin interactions in the relapse samples. Additionally, our analyses confirmed that a substantial proportion of open chromatin regions were lost upon disease relapse, highlighting the substantial chromatin remodeling in AML progression. Transcription factor enrichment analysis highlighted liver X receptors enriched in relapse-specific regions, which indicates that they could be potential therapeutic targets not only in chronic lymphocytic leukemia (Christopherson & Landay, 2009) but also in AML. Furthermore, differential gene expression analysis on longitudinal samples revealed that up-regulated genes are enriched in broad immune response pathways, including those related to cytokine signaling, especially interferon-alpha/beta signaling, while downregulated genes show enrichment in signaling pathways associated with Jak-STAT and interleukin.

Combined multi-omics data pinpointed three upregulated genes (*SDC2, NCAM2* and *IFI44*) that showed consistent effects on survival in a larger leukemia cohort. *TSPYL5* was downregulated in both of the relapse samples we tested, but its upregulation was associated with worse prognosis in the Leucegene cohort. This contradicts our observation and might be due to its differential response depending on the AML subtypes. Additionally, our analysis revealed that upregulation of genes in relapse samples could be attributed to both the formation of new active enhancer contacts and to the loss of chromatin interactions with silencer/poised enhancer regions upon relapse. In conclusion, our integrated analysis highlights distinct genomic and epigenomic profiles in relapse compared to diagnosis. Although the small sample size limits our study, these findings highlight the potential for further exploration in larger cohorts to elucidate the clinical relevance and therapeutic implications of these observations for KMT2Ar AML.

## Methods

### Patient samples

We received live-frozen mononuclear cells from bone marrow and/or peripheral blood of adult AML diagnosis and relapse paired samples from Bolzano General Hospital, Italy. Each sample contained >50% blast cells [Table 1]. The project was reviewed by the Hamad Bin Khalifa University Institutional Review Board and approved under protocol #QBRI-IRB 2020-02-017.

### Omni-C library preparation

We have employed the Omni-C™ kit, a sequence-independent endonuclease-based Dovetail Genomics™ proximity-ligation protocol. Briefly, 1 x 10^6^ live bone marrow cells were fixed in DSG (disuccinimidyl glutarate), a non-cleavable and membrane-permeable protein-protein crosslinker, followed by formaldehyde to reversibly crosslink in vivo DNA-protein interactions. The fixed cells were treated with DNaseI to digest chromatin. Next, for the proximity ligation, the chromatin ends were polished and biotin-tagged bridges were used to create chimeric molecules. The crosslink of lysate was reversed and the purified DNA was used for NGS library preparation. Finally, the library was enriched for ligation-containing chimeric molecules. The Omni-C libraries were sequenced on an average of 14x coverage on the Illumina HiSeq X Ten system with 151-base paired-end reads (>300M reads).

### ATAC-seq Library Preparation

One of our objectives was to interrogate active chromatin regions through an assessment of chromatin accessibility, which was evaluated using ATAC-seq, a method known for its capability to delineate regions of open chromatin. The Active Motif ATAC-seq kit was used to perform ATAC-seq on living cells in accordance with the Omni-ATAC-seq protocol described by (Corces et al., 2017). For ATAC-seq, cryopreserved bone marrow samples were slowly thawed using IMDM supplemented with 10% FBS and DNAse. Viability was calculated under a hemocytometer with trypan blue – samples with a viability <80% were subjected to a dead cell sort using MACS dead cell removal kit (cat: 130-090-101). 1x105 cells were taken forward for ATAC-seq.

Briefly, for sample preparation, 1L×L10^5^ cells were first pelleted and washed with ice-cold PBS. Subsequently, the cells were re-suspended in an ice-cold ATAC-Lysis buffer. Next, for tagmentation the isolated nuclei were incubated at 37°C for precisely 30 minutes while being shaken at 800 rpm in a transposition mixture containing 100 nM final transposase. For DNA purification Zymo DNA Clean and Concentrator-5 Kit was used. Following that, PCR amplifications of tagmented libraries were performed using 10 cycles of PCR and DNA was extracted using 60µl SPRI beads for size selection. To assess size distribution, PCR-amplified libraries were analyzed with Bioanalyzer. Finally, libraries were sequenced on the Illumina HiSeq 4000 platform to ∼50 million paired-end 100bp reads.

### RNA-seq library preparation

Total RNA was purified from 5X10^5^ live bone marrow cells using a Qiagen RNeasy Plus Micro kit. Briefly, the cells were disrupted and homogenized using RLT buffer. Genomic DNA eliminator spin columns were used to remove the DNA, and RNeasy MinElute spin columns were used to purify RNA. Total RNA-seq libraries were prepared using TruSeq Stranded Total RNA Library Prep Gold Kit. The prepared libraries were sequenced using the Illumina Nextseq platform to ∼60-50 million paired-end 101bp reads.

### Omni-C Data pre-processing

Omni-C libraries were processed through an in-house pipeline. The quality control (QC) of sequencing of Omni-C libraries was performed on fastq files using FastQC tool (Andrews et al., 2010). The adaptor sequences from the reads were trimmed using TrimGalore. The trimmed read pairs were aligned using BWA-MEM algorithm (version 0.7.17 or higher) to the GRCh38 version of the human genome. To find ligation junctions in Omni-C libraries, the “pairtools parse” module was used by setting MAPQ greater or equal to 40 and walks-policy as 5unique. The pairtools pipeline records the strand of each paired read and the outermost (5’) aligned base pair into a “pairsam” file upon identification of a ligation event in the alignment file. The pairsam format records Hi-C pair information along with SAM entries, which was then sorted using “pairtools sort”. The “pairtools dedup” command was used to remove PCR duplicates from sorted pairsam files and here we also produced the statistics of the library by using the flag “– output-stats”. Finally, the deduplicated pairsam files were then used to create two different files, such as pairs file and BAM file, which can be used for further downstream processing.

### Omni-C Library QC and Complexity

To check the QC of Omni-C libraries, we have used the stats files calculated by “pairtools dedup”, which contains information on total reads, mapped reads, duplicate reads, and total read pairs. In addition, we used the get_qc.py pipeline from Dovetail Genomics to summarize these stats in percentage and absolute values (Table). We have checked the complexity of Omni-C libraries using the lc_extrap utility of the preseq package from Smith lab (github.com/smithlabcode/preseq), which aims to predict the complexity of sequencing libraries.

### Contact Matrix

The contact maps, which are compressed and sparse formats are produced from the pairs files using Juicer tools (Durand et al., 2016). The pairs files were first converted into HiC files, which are highly compressed binary representations of the contact matrices using “pre” command of Juicer tools. The HiC contact matrices were finally visualized using Juicebox (J. T. Robinson et al., 2018).

### Differential Interaction Analysis

The systematic biases such as those arising from enzyme digestion, DNA ligation, and PCR amplification from Omni-C libraries were corrected using the HiCorr pipeline (Lu et al., 2020). To prepare the input, the BAM files were sorted according to co-ordinates and read pairs were then mapped to select the cis and trans read pairs. The HiCorr outputs were then used to apply a deep learning-based tool called DeepLoop (Zhang et al., 2022) to perform loop signal enhancement. We have used the pre-built models from DeepLoop to improve the sensitivity, robustness, and quantitation of Omni-C loops and output chromatin loop strength.

After that, we have created a count matrix (*M* _(i, j)_) where each row *(i)* represents a chromatin loop and column *(j)* represents a sample and it was populated with the loop strengths from DeepLoop. The count matrix was normalized using R Bioconductor package edgeR (M. D. Robinson et al., 2010). The normalized counts were used to calculate the standard deviation and we selected the top 100,000 most variable interactions based on standard deviation. Finally, the matrix with only the most variable interactions was used to perform differential interaction analysis using the R Bioconductor package limma (Ritchie et al., 2015).

### Omni-C Downstream Analysis

Exploratory analysis was performed in R version 4.3.1 using the Bioconductor package GenomicRanges (Lawrence et al., 2013). All interaction landscapes were visualized in WashU Epigenome Browser (Li et al., 2022).

### ATAC-seq Library Pre-processing

Briefly, all libraries underwent FastQC testing (Andrews et al., 2010) to evaluate the library quality and make sure that each library is free of significant problems like low read quality or adapter contamination. Next, we used Trimmomatic (Bolger et al., 2014) with default parameters to filter low-quality reads, and Truseq adaptors were trimmed off from the reads. Subsequently, these reads were aligned to the hg38 version of the human genome using Bowtie2 aligner, which created SAM files that were converted to BAM files using samtools. After that, mitochondrial, duplicate, and blacklisted reads were removed using samtools and bedtools. Reads were shifted to adjust for tn5 binding using the alignmentSieve tool. Finally, peaks were then called on the final processed BAM files using callpeak and BAMPE command of the MACS2 peak calling algorithm.

### ATAC-seq downstream analysis

MACS2 peaks were further used for downstream processing. Peaks were assigned to genomic elements using BioMart (Durinck et al., 2005) and TxDb.Hsapiens.UCSC.hg38.knownGene R Bioconductor packages (Carlson, et al., 2015). All exploratory analysis was performed in R version 4.3.1 using the GenomicRanges Bioconductor package (Lawrence et al., 2013).

### RNA-sequencing pre-processing

The QC of RNA-seq libraries was performed on fastq files using the FastQC tool (Andrews et al., 2010). The adaptor sequences from the reads were trimmed using TrimGalore. The trimmed reads were aligned to the hg38 genome using STAR and RSEM was used to calculate the expression values as expected counts from the aligned RNA-seq data. The count matrix was normalized using the R Bioconductor package edgeR (M. D. Robinson et al., 2010). Finally, the normalized counts were used to analyze differential gene expression using the R Bioconductor package limma (Ritchie et al., 2015). Significant up-regulated and down-regulated genes with adjusted p-value < 0.1 were selected based on log2 fold change >1 and log2 fold change < −1, respectively.

### RNA-seq downstream analysis

The normalized reads were used for all exploratory analysis and plots were generated using custom code in R version 4.3.1. Volcano plots were created using the R Bioconductor package ggplot2.

### Multi-Omics Data Integration

Significant differential Omni-C interactions were used to identify relapse-specific and diagnosis-specific anchors in each sample and subsequently, we specified diagnosis-specific ATAC-seq peaks by taking the overlap of two diagnosis samples and relapse-specific ATAC-seq peaks by taking the overlap of two relapse samples. Further, we removed the common peaks between diagnosis-specific and relapse-specific ATAC-seq peaks. Next, relapse-specific and diagnosis-specific Omni-C anchors and ATAC-seq peaks were overlapped using the Bioconductor package GenomicRanges (Lawrence et al., 2013) to investigate which differentially interacting regions are accessible. Finally, we assigned genes to these regions using BioMart (Durinck et al., 2005).

Furthermore, we investigated which differentially expressed genes are linked to differentially accessible DNA in differentially interacting regions (20Kb). Then, we calculated enrichment for differentially expressed genes in relapse and diagnosis-specific interactions in comparison with 10 random, size-matched sets of genes from the whole genome. Finally, the distal or other end of the interaction of upregulated genes that have diagnosis-specific Omni-C interactions and ATAC-seq peaks were overlapped with publicly available H3K27me3 ChIP-seq data (accession number GSM4565992) from CD34^+^ common myeloid progenitor cells, to test whether their upregulation can be attributed to the loss of a silences contact. For these analyses, the R Bioconductor package GenomicRanges (Lawrence et al., 2013) was used.

### Motif Enrichment

For the differential enrichment of transcription factor binding motifs within differentially accessible regions found in differentially interacting regions the Homer (Hypergeometric Optimization of Motif EnRichment) motif suite of tools was used (Heinz et al., 2010). The parameters findMotifsGenome.pl with -size -200,200 to centre peaks to a 400bp region and -bg to set background of peaks. Motifs with FDR<0.01 were considered significantly enriched.

Relapse-specific anchors and peaks were used as target sequences, and diagnosis-specific anchors and peaks were used as background sequences.

## Author Contributions

B.M. and N.A. conceived the study. C.C., S.D., and I.C. provided patient care and performed sample collection. N.A. performed the experiments, N.A. and W.V. performed data analysis, and N.A. wrote the manuscript. B.M. W.V., C.C., and S.D. reviewed and edited the manuscript, and all authors approved the manuscript.

## Data availability

All sequencing and processed data are uploaded to GEO accession GSE267375 and GSE267376.

## Acknowledgment

We are grateful to Roberto Gambato and Mirija Svaldi from the Hematology Laboratory in Bolzano Hospital for facilitating the patients’ samples collection. We would like to extend our thanks to Research Computing at TAMU-Q for using their high-performance computer (RAAD2).

---

**Supplementary Figure 1.** Schematic of data integration and WahsU arc plots. **A)** Schematic depicting the overlap of relapse-specific differential interactions with peaks and diagnosis-specific interactions and peaks. **B-F)** Arc plots from WashU Epigenome browser represent relapse-specific (B and C) and diagnosis-specific (D, E, and F) Omni-C interaction and ATAC-seq peaks of UBE2J1 and PARP8 as well as BMP8A, ST8SIA1, and HSPE2, respectively. The active or poised enhancers were validated using publicly available annotation data in browser, where the red track is for H3K27me3, blue is for H3K27ac, and green is for H3K4me3/1.

**Supplementary Figure 2.** Schematic and box plots. **A)** Bar plots showing the enrichment of biological processes and pathways based on significantly dysregulated genes between relapse versus diagnosis. **B)** Schematic portraying the interaction of promoter regions with distal poised enhancer marked by H3K27me3. **C)** Box plots representing down-regulated genes that overlapped with relapse-specific interactions and peaks and their comparison with the expression level of 10 different random sets of down-regulated genes. **D)** Box plots representing down-regulated genes that overlapped with diagnosis-specific interactions and peaks and their comparison with the expression level of 10 different random sets of down- regulated genes.

**Supplementary Figure 3.** **A-E)** Kaplan Meier plots showing the overall survival of patients from the Leucegene AML cohort based on gene expression data.

**Supplementary Table 1.** List of upregulated genes of relapse versus diagnosis AML samples with logFC >1 and adj-pval < 0.1.

**Supplementary Table 2.** List of downregulated genes of relapse versus diagnosis AML samples with logFC < -1 and adj-pval < 0.1.

## References

Ahn, J.-S., Kim, H.-J., Kim, Y.-K., Lee, S.-S., Ahn, S.-Y., Jung, S.-H., Yang, D.-H., Lee, J.-J., Park, H. J., Lee, J.-Y., Choi, S. H., Jung, C. W., Jang, J.-H., Kim, H. J., Moon, J. H., Sohn, S. K., Lee, Y. J., Won, J.-H., Kim, S.-H., … Kim, D. D. H. (2018). Assessment of a new genomic classification system in acute myeloid leukemia with a normal karyotype. Oncotarget, 9(4), 4961–4968. 10.18632/oncotarget.23575

Akl, M. R., Nagpal, P., Ayoub, N. M., Prabhu, S. A., Gliksman, M., Tai, B., Hatipoglu, A., Goy, A., & Suh, K. S. (2015). Molecular and clinical profiles of syndecan-1 in solid and hematological cancer for prognosis and precision medicine. Oncotarget, 6(30), 28693– 28715. https://www.ncbi.nlm.nih.gov/pmc/articles/PMC4745686/

Andrews, S., Biggins, L., Inglesfield, S., Carr, H., & Montgomery, J. (2010). *FastQC A Quality Control tool for High Throughput Sequence Data* [Computer software]. https://www.bioinformatics.babraham.ac.uk/projects/fastqc/

Bejanyan, N., Weisdorf, D. J., Logan, B. R., Wang, H.-L., Devine, S. M., de Lima, M., Bunjes, D. W., & Zhang, M.-J. (2015). Survival of patients with acute myeloid leukemia relapsing after allogeneic hematopoietic cell transplantation: A center for international blood and marrow transplant research study. Biology of Blood and Marrow Transplantation: Journal of the American Society for Blood and Marrow Transplantation, 21(3), 454–459. 10.1016/j.bbmt.2014.11.007

Bolger, A. M., Lohse, M., & Usadel, B. (2014). Trimmomatic: A flexible trimmer for Illumina sequence data. Bioinformatics, 30(15), 2114–2120. 10.1093/bioinformatics/btu170

Canarte, V., Moser-Katz, T., Gavile, C., Barwick, B. G., Lee, K. P., & Boise, L. H. (2023). SDC2 Expression Is Increased in Myeloma Cells in Response to Loss of Pro-Survival Surface Proteins, CD28 and CD86. Blood, 142(Supplement 1), 3299. 10.1182/blood-2023-190453

Christopherson, K. W., II, & Landay, A. (2009). Editorial: Liver X receptor α (LXRα) as a therapeutic target in chronic lymphocytic leukemia (CLL). Journal of Leukocyte Biology, 86(5), 1019–1021. 10.1189/jlb.0509295

Corces, M. R., Trevino, A. E., Hamilton, E. G., Greenside, P. G., Sinnott-Armstrong, N. A., Vesuna, S., Satpathy, A. T., Rubin, A. J., Montine, K. S., Wu, B., Kathiria, A., Cho, S. W., Mumbach, M. R., Carter, A. C., Kasowski, M., Orloff, L. A., Risca, V. I., Kundaje, A., Khavari, P. A., … Chang, H. Y. (2017). An improved ATAC-seq protocol reduces background and enables interrogation of frozen tissues. Nature Methods, 14(10), 959– 962. 10.1038/nmeth.4396

DiNardo, C. D., & Cortes, J. E. (2016). Mutations in AML: Prognostic and therapeutic implications. Hematology: The American Society of Hematology Education Program, 2016(1), 348–355. https://www.ncbi.nlm.nih.gov/pmc/articles/PMC6142505/

Döhner, H., Estey, E., Grimwade, D., Amadori, S., Appelbaum, F. R., Büchner, T., Dombret, H., Ebert, B. L., Fenaux, P., Larson, R. A., Levine, R. L., Lo-Coco, F., Naoe, T., Niederwieser, D., Ossenkoppele, G. J., Sanz, M., Sierra, J., Tallman, M. S., Tien, H.-F., … Bloomfield, C. D. (2017). Diagnosis and management of AML in adults: 2017 ELN recommendations from an international expert panel. Blood, 129(4), 424–447. 10.1182/blood-2016-08-733196

Döhner, H., Wei, A. H., Appelbaum, F. R., Craddock, C., DiNardo, C. D., Dombret, H., Ebert, B. L., Fenaux, P., Godley, L. A., Hasserjian, R. P., Larson, R. A., Levine, R. L., Miyazaki, Y., Niederwieser, D., Ossenkoppele, G., Röllig, C., Sierra, J., Stein, E. M., Tallman, M. S., … Löwenberg, B. (2022). Diagnosis and management of AML in adults: 2022 recommendations from an international expert panel on behalf of the ELN. Blood, 140(12), 1345–1377. 10.1182/blood.2022016867

Döhner, H., Weisdorf, D. J., & Bloomfield, C. D. (2015). Acute Myeloid Leukemia. New England Journal of Medicine, 373(12), 1136–1152. 10.1056/NEJMra1406184

Durand, N. C., Shamim, M. S., Machol, I., Rao, S. S. P., Huntley, M. H., Lander, E. S., & Aiden, E. L. (2016). Juicer Provides a One-Click System for Analyzing Loop-Resolution Hi-C Experiments. Cell Systems, 3(1), 95–98. 10.1016/j.cels.2016.07.002

Durinck, S., Moreau, Y., Kasprzyk, A., Davis, S., De Moor, B., Brazma, A., & Huber, W. (2005). BioMart and Bioconductor: A powerful link between biological databases and microarray data analysis. *Bioinformatics (Oxford*, England), 21(16), 3439–3440. 10.1093/bioinformatics/bti525

Farrar, J. E., Schuback, H. L., Ries, R. E., Wai, D., Hampton, O. A., Trevino, L. R., Alonzo, T. A., Guidry Auvil, J. M., Davidsen, T. M., Gesuwan, P., Hermida, L., Muzny, D. M., Dewal, N., Rustagi, N., Lewis, L. R., Gamis, A. S., Wheeler, D. A., Smith, M. A., Gerhard, D. S., & Meshinchi, S. (2016). Genomic Profiling of Pediatric Acute Myeloid Leukemia Reveals a Changing Mutational Landscape from Disease Diagnosis to Relapse. Cancer Research, 76(8), 2197–2205. 10.1158/0008-5472.CAN-15-1015

Gröschel, S., Sanders, M. A., Hoogenboezem, R., Zeilemaker, A., Havermans, M., Erpelinck, C., Bindels, E. M. J., Beverloo, H. B., Döhner, H., Löwenberg, B., Döhner, K., Delwel, R., & Valk, P. J. M. (2015). Mutational spectrum of myeloid malignancies with inv(3)/t(3;3) reveals a predominant involvement of RAS/RTK signaling pathways. Blood, 125(1), 133–139. 10.1182/blood-2014-07-591461

Howlader N, Noone AM, Krapcho M, Miller D, Brest A, Yu M, Ruhl J, Tatalovich Z, Mariotto A, Lewis DR, Chen HS, Feuer EJ, & Cronin KA (eds). (2021). SEER Cancer Statistics Review, 1975-2018, *National Cancer Institute*. Bethesda, MD, https://seer.cancer.gov/csr/1975_2018/, based on November 2020 SEER data submission, posted to the SEER web site, April 2021. SEER. https://seer.cancer.gov/csr/1975_2018/index.html

Issa, G. C., Aldoss, I., DiPersio, J., Cuglievan, B., Stone, R., Arellano, M., Thirman, M. J., Patel, M. R., Dickens, D. S., Shenoy, S., Shukla, N., Kantarjian, H., Armstrong, S. A., Perner, F., Perry, J. A., Rosen, G., Bagley, R. G., Meyers, M. L., Ordentlich, P., … Stein, E. M. (2023). The menin inhibitor revumenib in KMT2A-rearranged or NPM1-mutant leukaemia. Nature, 615(7954), Article 7954. 10.1038/s41586-023-05812-3

Jan, M., Snyder, T. M., Corces-Zimmerman, M. R., Vyas, P., Weissman, I. L., Quake, S. R., & Majeti, R. (2012). Clonal evolution of preleukemic hematopoietic stem cells precedes human acute myeloid leukemia. Science Translational Medicine, 4(149), 149–118. 10.1126/scitranslmed.3004315

Karlsson, L., Forestier, E., Hasle, H., Jahnukainen, K., Jónsson, Ó. G., Lausen, B., Norén Nyström, U., Palle, J., Tierens, A., Zeller, B., & Abrahamsson, J. (2017). Outcome after intensive reinduction therapy and allogeneic stem cell transplant in paediatric relapsed acute myeloid leukaemia. British Journal of Haematology, 178(4), 592–602. 10.1111/bjh.14720

Kerbs, P., Vosberg, S., Krebs, S., Graf, A., Blum, H., Swoboda, A., Batcha, A. M. N., Mansmann, U., Metzler, D., Heckman, C. A., Herold, T., & Greif, P. A. (2022). Fusion gene detection by RNA-sequencing complements diagnostics of acute myeloid leukemia and identifies recurring *NRIP1-MIR99AHG* rearrangements. Haematologica, 107(1), Article 1. 10.3324/haematol.2021.278436

Kern, W., Haferlach, T., Schnittger, S., Ludwig, W. D., Hiddemann, W., & Schoch, C. (2002). Karyotype instability between diagnosis and relapse in 117 patients with acute myeloid leukemia: Implications for resistance against therapy. Leukemia, 16(10), 2084–2091. 10.1038/sj.leu.2402654

Kishtagari, A., & Levine, R. L. (2021). The Role of Somatic Mutations in Acute Myeloid Leukemia Pathogenesis. Cold Spring Harbor Perspectives in Medicine, 11(4), a034975. 10.1101/cshperspect.a034975

Krivtsov, A. V., & Armstrong, S. A. (2007). MLL translocations, histone modifications and leukaemia stem-cell development. Nature Reviews Cancer, 7(11), Article 11. 10.1038/nrc2253

Lambo, S., Trinh, D. L., Ries, R. E., Jin, D., Setiadi, A., Ng, M., Leblanc, V. G., Loken, M. R., Brodersen, L. E., Dai, F., Pardo, L. M., Ma, X., Vercauteren, S. M., Meshinchi, S., & Marra, M. A. (2023). A longitudinal single-cell atlas of treatment response in pediatric AML. Cancer Cell, 41(12), 2117–2135.e12. 10.1016/j.ccell.2023.10.008

Lawrence, M., Huber, W., Pagès, H., Aboyoun, P., Carlson, M., Gentleman, R., Morgan, M. T., & Carey, V. J. (2013). Software for Computing and Annotating Genomic Ranges. PLOS Computational Biology, 9(8), e1003118. 10.1371/journal.pcbi.1003118

Levin, M., Stark, M., Ofran, Y., & Assaraf, Y. G. (2021). Deciphering molecular mechanisms underlying chemoresistance in relapsed AML patients: Towards precision medicine overcoming drug resistance. Cancer Cell International, 21(1), 53. 10.1186/s12935-021-01746-w

Li, D., Purushotham, D., Harrison, J. K., Hsu, S., Zhuo, X., Fan, C., Liu, S., Xu, V., Chen, S., Xu, J., Ouyang, S., Wu, A. S., & Wang, T. (2022). WashU Epigenome Browser update 2022. Nucleic Acids Research, 50(W1), W774–W781. 10.1093/nar/gkac238

Liedtke, M., & Cleary, M. L. (2009). Therapeutic targeting of MLL. Blood, 113(24), 6061–6068. 10.1182/blood-2008-12-197061

Lu, L., Liu, X., Huang, W.-K., Giusti-Rodríguez, P., Cui, J., Zhang, S., Xu, W., Wen, Z., Ma, S., Rosen, J. D., Xu, Z., Bartels, C. F., Kawaguchi, R., Hu, M., Scacheri, P. C., Rong, Z., Li, Y., Sullivan, P. F., Song, H., … Jin, F. (2020). Robust Hi-C Maps of Enhancer-Promoter Interactions Reveal the Function of Non-coding Genome in Neural Development and Diseases. Molecular Cell, 79(3), 521–534.e15. 10.1016/j.molcel.2020.06.007

Masetti, R., Castelli, I., Astolfi, A., Bertuccio, S. N., Indio, V., Togni, M., Belotti, T., Serravalle, S., Tarantino, G., Zecca, M., Pigazzi, M., Basso, G., Pession, A., & Locatelli, F. (2016). Genomic complexity and dynamics of clonal evolution in childhood acute myeloid leukemia studied with whole-exome sequencing. Oncotarget, 7(35), 56746–56757. 10.18632/oncotarget.10778

Metzeler, K. H., Herold, T., Rothenberg-Thurley, M., Amler, S., Sauerland, M. C., Görlich, D., Schneider, S., Konstandin, N. P., Dufour, A., Bräundl, K., Ksienzyk, B., Zellmeier, E., Hartmann, L., Greif, P. A., Fiegl, M., Subklewe, M., Bohlander, S. K., Krug, U., Faldum, A., … on behalf of the AMLCG Study Group. (2016). Spectrum and prognostic relevance of driver gene mutations in acute myeloid leukemia. Blood, 128(5), 686–698. 10.1182/blood-2016-01-693879

Meyer, C., Burmeister, T., Gröger, D., Tsaur, G., Fechina, L., Renneville, A., Sutton, R., Venn, N. C., Emerenciano, M., Pombo-de-Oliveira, M. S., Barbieri Blunck, C., Almeida Lopes, B., Zuna, J., Trka, J., Ballerini, P., Lapillonne, H., De Braekeleer, M., Cazzaniga, G., Corral Abascal, L., … Marschalek, R. (2018). The MLL recombinome of acute leukemias in 2017. Leukemia, 32(2), Article 2. 10.1038/leu.2017.213

Northcott, P. A., Lee, C., Zichner, T., Stütz, A. M., Erkek, S., Kawauchi, D., Shih, D. J. H., Hovestadt, V., Zapatka, M., Sturm, D., Jones, D. T. W., Kool, M., Remke, M., Cavalli, F. M. G., Zuyderduyn, S., Bader, G. D., VandenBerg, S., Esparza, L. A., Ryzhova, M., … Pfister, S. M. (2014). Enhancer hijacking activates GFI1 family oncogenes in medulloblastoma. Nature, 511(7510), Article 7510. 10.1038/nature13379

Papaemmanuil, E., Gerstung, M., Bullinger, L., Gaidzik, V. I., Paschka, P., Roberts, N. D., Potter, N. E., Heuser, M., Thol, F., Bolli, N., Gundem, G., Van Loo, P., Martincorena, I., Ganly, P., Mudie, L., McLaren, S., O’Meara, S., Raine, K., Jones, D. R., … Campbell, P. J. (2016). Genomic Classification and Prognosis in Acute Myeloid Leukemia. New England Journal of Medicine, 374(23), 2209–2221. 10.1056/NEJMoa1516192

Rasche, M., Zimmermann, M., Steidel, E., Alonzo, T., Aplenc, R., Bourquin, J.-P., Boztug, H., Cooper, T., Gamis, A. S., Gerbing, R. B., Janotova, I., Klusmann, J.-H., Lehrnbecher, T., Mühlegger, N., Neuhoff, N. v, Niktoreh, N., Sramkova, L., Stary, J., Waack, K., … Reinhardt, D. (2021). Survival Following Relapse in Children with Acute Myeloid Leukemia: A Report from AML-BFM and COG. Cancers, 13(10), Article 10. 10.3390/cancers13102336

Ritchie, M. E., Phipson, B., Wu, D., Hu, Y., Law, C. W., Shi, W., & Smyth, G. K. (2015). Limma powers differential expression analyses for RNA-sequencing and microarray studies. Nucleic Acids Research, 43(7), e47. 10.1093/nar/gkv007

Robinson, J. T., Turner, D., Durand, N. C., Thorvaldsdóttir, H., Mesirov, J. P., & Aiden, E. L. (2018). Juicebox.js Provides a Cloud-Based Visualization System for Hi-C Data. Cell Systems, 6(2), 256–258.e1. 10.1016/j.cels.2018.01.001

Robinson, M. D., McCarthy, D. J., & Smyth, G. K. (2010). edgeR: A Bioconductor package for differential expression analysis of digital gene expression data. Bioinformatics, 26(1), 139–140. 10.1093/bioinformatics/btp616

Schlenk, R. F., Jaramillo, S., & Müller-Tidow, C. (2018). Improving consolidation therapy in acute myeloid leukemia—A tough nut to crack. Haematologica, 103(10), 1579–1581. 10.3324/haematol.2018.200485

Schwenger, E., & Steidl, U. (2021). An Evolutionary Approach to Clonally Complex Hematologic Disorders. Blood Cancer Discovery, 2(3), 201–215. 10.1158/2643-3230.BCD-20-0219

Siegel, R. L., Miller, K. D., Wagle, N. S., & Jemal, A. (2023). Cancer statistics, 2023. CA: A Cancer Journal for Clinicians, 73(1), 17–48. 10.3322/caac.21763

Stein, E. M., DiNardo, C. D., Pollyea, D. A., Fathi, A. T., Roboz, G. J., Altman, J. K., Stone, R. M., DeAngelo, D. J., Levine, R. L., Flinn, I. W., Kantarjian, H. M., Collins, R., Patel, M. R., Frankel, A. E., Stein, A., Sekeres, M. A., Swords, R. T., Medeiros, B. C., Willekens, C., … Tallman, M. S. (2017). Enasidenib in mutant IDH2 relapsed or refractory acute myeloid leukemia. Blood, 130(6), 722–731. 10.1182/blood-2017-04-779405

Stone, R. M., Mandrekar, S. J., Sanford, B. L., Laumann, K., Geyer, S., Bloomfield, C. D., Thiede, C., Prior, T. W., Döhner, K., Marcucci, G., Lo-Coco, F., Klisovic, R. B., Wei, A., Sierra, J., Sanz, M. A., Brandwein, J. M., de Witte, T., Niederwieser, D., Appelbaum, F. R., … Döhner, H. (2017). Midostaurin plus Chemotherapy for Acute Myeloid Leukemia with a FLT3 Mutation. New England Journal of Medicine, 377(5), 454–464. 10.1056/NEJMoa1614359

Tirtakusuma, R., Szoltysek, K., Milne, P., Grinev, V. V., Ptasinska, A., Chin, P. S., Meyer, C., Nakjang, S., Hehir-Kwa, J. Y., Williamson, D., Cauchy, P., Keane, P., Assi, S. A., Ashtiani, M., Kellaway, S. G., Imperato, M. R., Vogiatzi, F., Schweighart, E. K., Lin, S.,… Bomken, S. (2022). Epigenetic regulator genes direct lineage switching in MLL/AF4 leukemia. Blood, 140(17), 1875–1890. 10.1182/blood.2021015036

van der Sluis, I. M., de Lorenzo, P., Kotecha, R. S., Attarbaschi, A., Escherich, G., Nysom, K., Stary, J., Ferster, A., Brethon, B., Locatelli, F., Schrappe, M., Scholte-van Houtem, P. E., Valsecchi, M. G., & Pieters, R. (2023). Blinatumomab Added to Chemotherapy in Infant Lymphoblastic Leukemia. New England Journal of Medicine, 388(17), 1572–1581. 10.1056/NEJMoa2214171

Verma, D., Kantarjian, H., Faderl, S., O’Brien, S., Pierce, S., Vu, K., Freireich, E., Keating, M., Cortes, J., & Ravandi, F. (2010). Late relapses in acute myeloid leukemia: Analysis of characteristics and outcome. Leukemia & Lymphoma, 51(5), 778–782. 10.3109/10428191003661852

Vicente-Dueñas, C., Hauer, J., Cobaleda, C., Borkhardt, A., & Sánchez-García, I. (2018). Epigenetic Priming in Cancer Initiation. Trends in Cancer, 4(6), 408–417. 10.1016/j.trecan.2018.04.007

Wang, B., Liu, Y., Hou, G., Wang, L., Lv, N., Xu, Y., Xu, Y., Wang, X., Xuan, Z., Jing, Y., Li, H., Jin, X., Deng, A., Wang, L., Gao, X., Dou, L., Liang, J., Chen, C., Li, Y., & Yu, L. (2016). Mutational spectrum and risk stratification of intermediate-risk acute myeloid leukemia patients based on next-generation sequencing. Oncotarget, 7(22), 32065–32078. 10.18632/oncotarget.7028

Zhang, S., Plummer, D., Lu, L., Cui, J., Xu, W., Wang, M., Liu, X., Prabhakar, N., Shrinet, J., Srinivasan, D., Fraser, P., Li, Y., Li, J., & Jin, F. (2022). DeepLoop robustly maps chromatin interactions from sparse allele-resolved or single-cell Hi-C data at kilobase resolution. Nature Genetics, 54(7), Article 7. 10.1038/s41588-022-01116-w

Zhu, H., Uusküla-Reimand, L., Isaev, K., Wadi, L., Alizada, A., Shuai, S., Huang, V., Aduluso- Nwaobasi, D., Paczkowska, M., Abd-Rabbo, D., Ocsenas, O., Liang, M., Thompson, J. D., Li, Y., Ruan, L., Krassowski, M., Dzneladze, I., Simpson, J. T., Lupien, M., … Reimand, J. (2020). Candidate Cancer Driver Mutations in Distal Regulatory Elements and Long-Range Chromatin Interaction Networks. Molecular Cell, 77(6), 1307–1321.e10. 10.1016/j.molcel.2019.12.027

