## Supplementary figures and images for "Multi-omics analysis of longitudinal patient samples reveals the molecular mechanism of AML progression"

### Supplementary Figure 1

Supplementary Figure 1

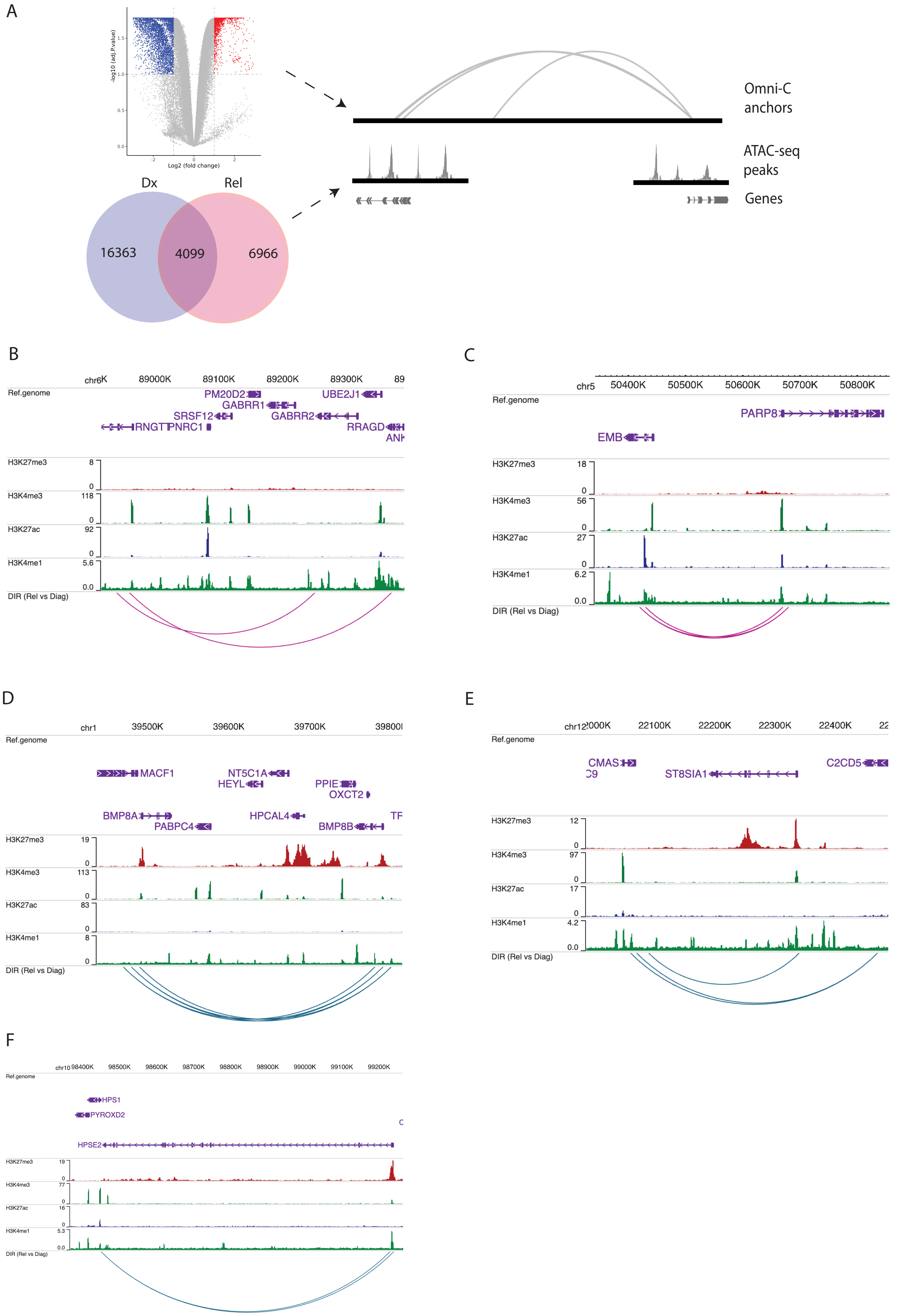

### Supplementary Figure 2

Supplimentary Figure 2

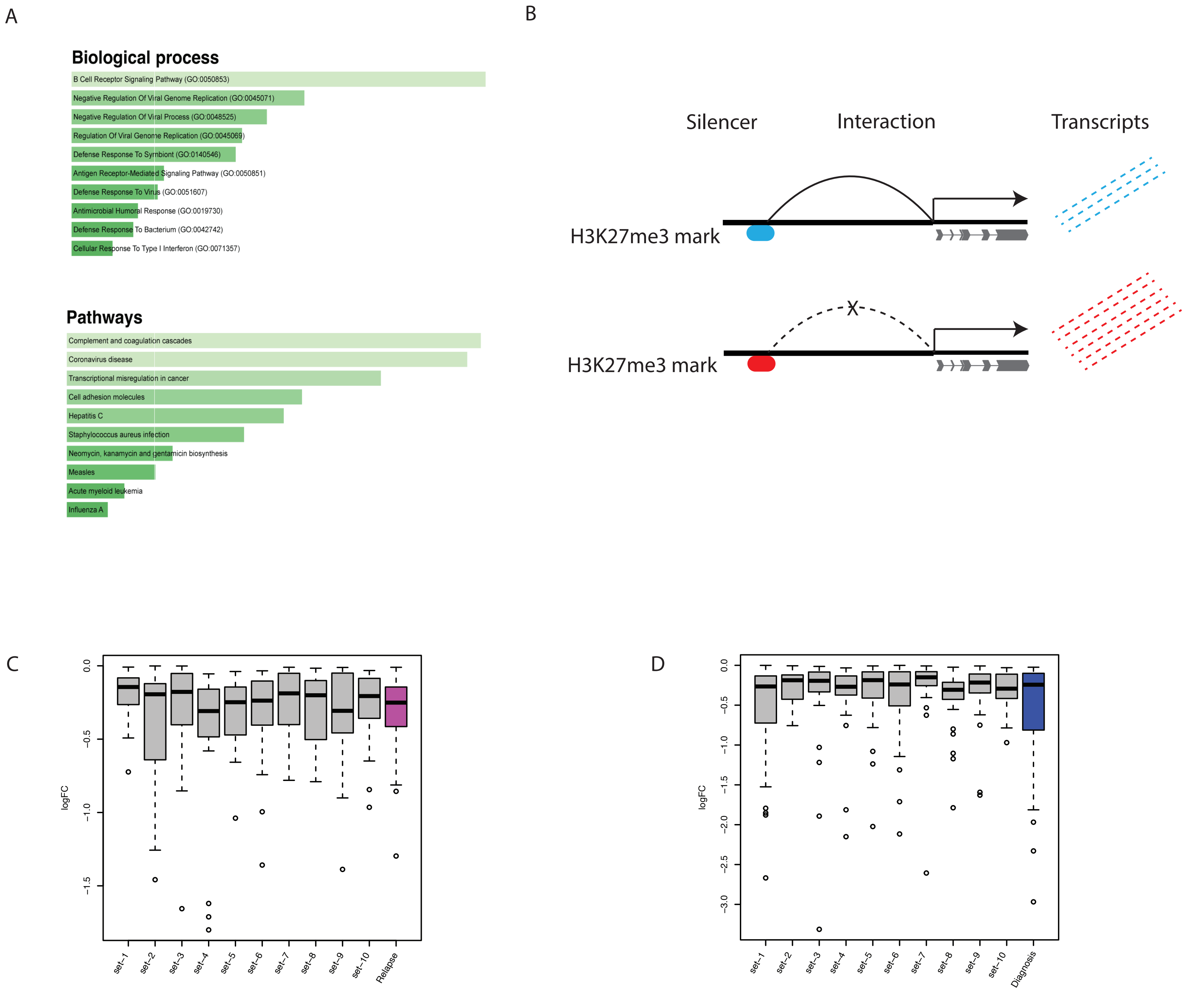
