## Supplementary Figure 3 for "Multi-omics analysis of longitudinal patient samples reveals the molecular mechanism of AML progression"

Supplimentary Figure 3

A

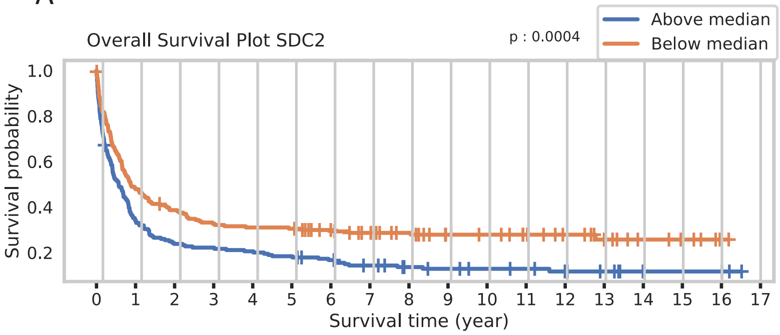


Numbers at risk

| (Year) | 0 | 1 | 2 | 3 | 4 | 5 | 6 | 7 | 8 | 9 | 10 | 11 | 12 | 13 | 14 | 15 | 16 | 17 |
|---|---|---|---|---|---|---|---|---|---|---|---|---|---|---|---|---|---|---|
| Above median: | 186 | 63 | 44 | 41 | 38 | 34 | 29 | 23 | 18 | 15 | 13 | 12 | 9 | 7 | 3 | 3 | 3 | 0 |
| Below median: | 187 | 89 | 71 | 60 | 57 | 56 | 48 | 41 | 36 | 29 | 27 | 23 | 16 | 11 | 7 | 5 | 1 | 0 |

B

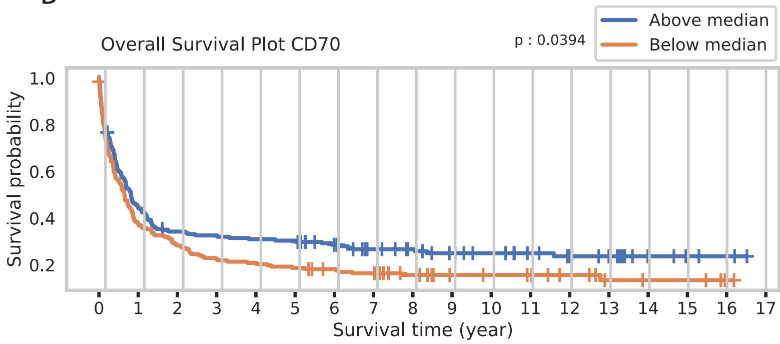


Numbers at risk

| (Year) | 0 | 1 | 2 | 3 | 4 | 5 | 6 | 7 | 8 | 9 | 10 | 11 | 12 | 13 | 14 | 15 | 16 | 17 |
|---|---|---|---|---|---|---|---|---|---|---|---|---|---|---|---|---|---|---|
| Above median: | 185 | 82 | 62 | 59 | 56 | 55 | 46 | 36 | 32 | 27 | 24 | 20 | 16 | 13 | 6 | 4 | 3 | 0 |
| Below median: | 188 | 70 | 53 | 42 | 39 | 35 | 31 | 28 | 22 | 17 | 16 | 15 | 9 | 5 | 4 | 4 | 1 | 0 |

C

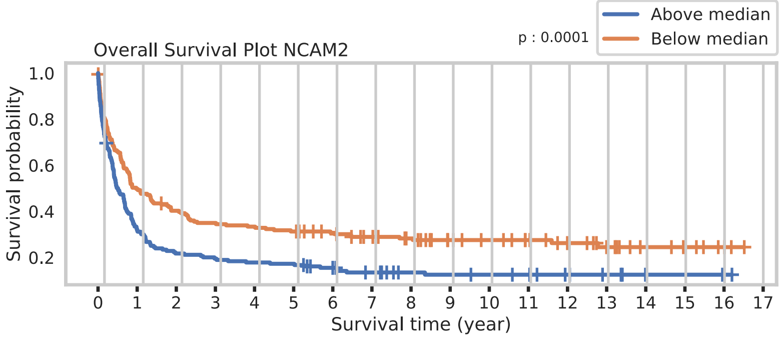


Numbers at risk

| (Year) | 0 | 1 | 2 | 3 | 4 | 5 | 6 | 7 | 8 | 9 | 10 | 11 | 12 | 13 | 14 | 15 | 16 | 17 |
|---|---|---|---|---|---|---|---|---|---|---|---|---|---|---|---|---|---|---|
| Below median: | 192 | 95 | 76 | 66 | 63 | 59 | 53 | 45 | 40 | 31 | 28 | 24 | 18 | 13 | 8 | 6 | 3 | 0 |
| Above median: | 181 | 57 | 39 | 35 | 32 | 31 | 24 | 19 | 14 | 13 | 12 | 11 | 7 | 5 | 2 | 2 | 1 | 0 |

D

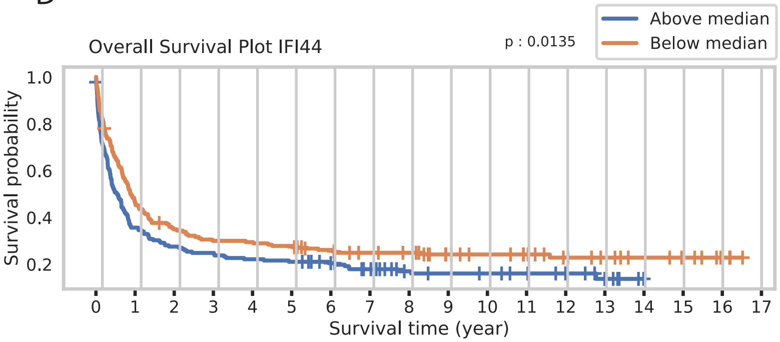


Numbers at risk

| (Year) | 0 | 1 | 2 | 3 | 4 | 5 | 6 | 7 | 8 | 9 | 10 | 11 | 12 | 13 | 14 | 15 | 16 | 17 |
|---|---|---|---|---|---|---|---|---|---|---|---|---|---|---|---|---|---|---|
| Above median: | 186 | 66 | 51 | 46 | 41 | 39 | 33 | 26 | 18 | 16 | 15 | 13 | 9 | 5 | 0 | 0 | 0 | 0 |
| Below median: | 187 | 86 | 64 | 55 | 54 | 51 | 44 | 38 | 36 | 28 | 25 | 22 | 16 | 13 | 10 | 8 | 4 | 0 |

E

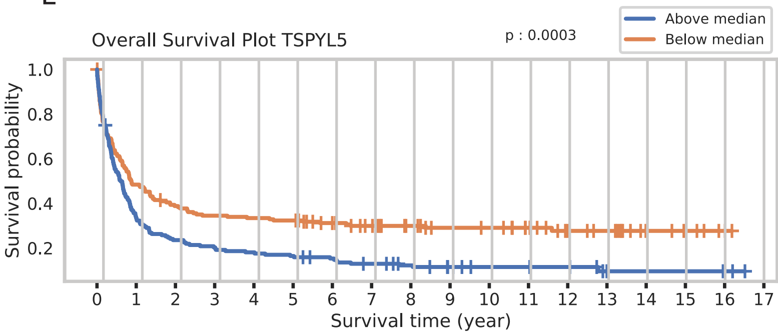


Numbers at risk

| (Year) | 0 | 1 | 2 | 3 | 4 | 5 | 6 | 7 | 8 | 9 | 10 | 11 | 12 | 13 | 14 | 15 | 16 | 17 |
|---|---|---|---|---|---|---|---|---|---|---|---|---|---|---|---|---|---|---|
| Below median: | 189 | 91 | 72 | 64 | 62 | 60 | 51 | 43 | 37 | 32 | 31 | 26 | 17 | 15 | 7 | 5 | 2 | 0 |
| Above median: | 184 | 61 | 43 | 37 | 33 | 30 | 26 | 21 | 17 | 12 | 9 | 9 | 8 | 3 | 3 | 3 | 2 | 0 |
